# Comparative genomics suggests limited variability and similar evolutionary patterns between major clades of SARS-CoV-2

**DOI:** 10.1101/2020.03.30.016790

**Authors:** Matteo Chiara, David S. Horner, Carmela Gissi, Graziano Pesole

**Affiliations:** Department of Biosciences, University of Milan, Italy; Institute of Biomembranes, Bioenergetics and Molecular Biotechnologies, Consiglio Nazionale delle Ricerche, Bari, Italy; Department of Biosciences, Biotechnology and Biopharmaceutics, University of Bari “A. Moro, Italy

## Abstract

Phylogenomic analysis of SARS-CoV-2 as available from publicly available repositories suggests the presence of 3 prevalent groups of viral episomes (super-clades), which are mostly associated with outbreaks in distinct geographic locations (China, USA and Europe). While levels of genomic variability between SARS-CoV-2 isolates are limited, to our knowledge, it is not clear whether the observed patterns of variability in viral super-clades reflect ongoing adaptation of SARS-CoV-2, or merely genetic drift and founder effects. Here, we analyze more than 1100 complete, high quality SARS-CoV-2 genome sequences, and provide evidence for the absence of distinct evolutionary patterns/signatures in the genomes of the currently known major clades of SARS-CoV-2. Our analyses suggest that the presence of distinct viral episomes at different geographic locations are consistent with founder effects, coupled with the rapid spread of this novel virus. We observe that while cross species adaptation of the virus is associated with hypervariability of specific protein coding regions (including the RDB domain of the spike protein), the more variable genomic regions between extant SARS-CoV-2 episomes correspond with the 3’ and 5’ UTRs, suggesting that at present viral protein coding genes should not be subjected to different adaptive evolutionary pressures in different viral strains. Although this study can not be conclusive, we believe that the evidence presented here is strongly consistent with the notion that the biased geographic distribution of SARS-CoV-2 isolates should not be associated with adaptive evolution of this novel pathogen.

## Introduction

The SARS-CoV-2 pandemic (Poon and Peiris, 2020) poses the greatest global health and socioeconomic threat since the second world war. Complete genomic sequences of viral isolates from diverse geographic sites, have rapidly been made available through dedicated resources (Shu and McCauley, 2017, Goodacre et al,2018) facilitating comparative genomics studies, identification of putative therapeutic targets (Zhou et al, 2020, Chen et al 2020, Robson 2020) and the development of effective prevention and monitoring strategies (Qiang et al 2020). Analyses of available genomic sequences, according to GISAID EpiCoV, suggest major viral clades, S, V and G which, collectively, circumscribe more than 69% of the characterised isolates. Strikingly, these clades show a markedly biased prevalence in different areas, with the S clade accounting for more than 72% of the viral isolates characterized in the USA, and the G clade comprising more than 74% of those that have been sequenced in Europe.

Importantly, while the G-clade was initially considered to be composed of viral strains which were not observed in China, availability of additional genomic sequences suggests that this clade as well should be nested within isolates from Shanghai. Although comparative analyses suggest that genomic variability between different isolates of SARS-CoV-2 is generally low (Lu et al, 2020, Zhang e t al, 2020, Tang et al, 2020), the fact that distinct viral episomes show a highly biased geographic distribution is potentially alarming, as, at present, it is not completely clear whether frequent variants reflect the adaptive processes, which result in the emergence of novel, and potentially more virulent strains. Moreover at present it is unclear whether the genomic variability of major clades of SARS-CoV-2 and their biased geographic distribution, could explain-at least in part-apparent differing rates of lethality observed worldwide (Baud et al 2020).

In the present study, exploiting curated viral genomic sequences, we present analyses of more than 1100 complete SARS-CoV-2 genomes, identified from 5 continents and more than 45 countries. By contrasting evolutionary patterns associated with the most prevalent viral clades, with those observed between closely related viral strains isolated from various species, we provide insights into the evolutionary dynamics and adaptation of SARS-CoV-2 like viruses to different hosts and the evolutionary patterns of the major clades of SARS-CoV-2.

We show that while the majority of the genomic variants that discriminate between major viral clades cause non synonymous substitutions in protein coding genes, including genes implicated in the modulation of the virulence of SARS-CoV-2 such as the spike protein and the RNA dependent polymerase (Weiss and Navas-Martin, 2005), the major clades of SARS-CoV-2 show nearly identical patterns of genomic variability as well as the absence of signatures that are normally associated with adaptive evolution at protein coding loci. Indeed, the major clades of SARS-CoV-2 are identified only by a limited number of clade-specific genetic variants and show very modest variability. Notably, variable sites are enriched in the 5’ and 3’ non coding regions of the genome, unlike genomic sites which are hyper-variable between closely related strains with distinct host specificites.

While our observations cannot be considered conclusive, we believe that the available data are broadly consistent with the notion that the biased geographic distribution of SARS-CoV-2 isolates is not associated with adaptive evolution of this novel pathogen, but rather with extensive founder effects coupled with the rapid spread of this pathogen in diverse geographic zones.

## Results

A total of 1113 complete, high quality SARS-CoV-2 genomic sequences, as well as of 2 SARS-CoV-2 like viruses isolated from non-human hosts (bats and pangolins (Zhou et al 2020, Matthew et al 2020), were retrieved from the GISAID EpiCoV portal on March 24th 2020. Associated metadata (Supplementary Table S1) show that these isolates included in cover more than 45 countries in 5 continents. As expected, the geographic distribution of the data closely reflects geographic prevalence of the pandemic, although notable exceptions include limited public data from Italy, one of the early hotspots of the pandemic.

Genomic sequences obtained from GISAID EpiCoV were aligned to the reference SARS-CoV-2 assembly (Refseq (O’Leary et al, 2016) accession NC_045512.2), i.e., the presumably ancestral Wuhan isolate (Wu et al, 2020), and derive a phenetic matrix of presence/absence of the variants. Substitution patterns of nucleotide residues, as shown in Supplementary Table S2 and Supplementary Figure S1, show a clear prevalence of C->T substitutions with respect to the presumably ancestral Wuhan isolate, with C->T representing 38% of all the observed distinct variants and an almost 4 fold enrichment of C->T with respect to T->C. Strikingly, the same pattern is not recovered when the SARS-CoV-2 genome is compared with genomic sequences of closely related viral specimens isolated from non-human hosts, suggesting a specific error/substitution pattern of the SARS-CoV-2 RNA dependent RNA polymerase. This observation is confirmed even when only polymorphic sites common between 2 or more genomes are considered. Intriguingly, analyses of the substitution patterns of the coronavirus associated with the 2003-2004 SARS outbreak (Chinese SARS Molecular Epidemiology Consortium, 2004, Song et al 2005) do show a similar, but less marked tendency of increased C->T substitutions.

Clustering of viral episomes based on 844 genetic variants present in at least 2 genomes - Fig 1 (see Materials and methods)-delineates 3 super-clades of viral strains, consistent with the classification of the isolates proposed by the GISAID EpiCoV portal (Shu and McCauley, 2017). Super-clade I is a superset of clade V as defined in GISAID, and contains viral sequences that have a limited variability with respect to the reference genome and were isolated in Europe and Asia. Super-clade II corresponds with the S-clade as identified by GISAID and incorporates viral isolates which are prevalently from the USA and from China. Finally Super-clade III corresponds with the G-clade as defined by GISAID and is formed mostly by European sequences. Therefore, a highly biased geographic distribution (Figure 1), consistent with previous reports, is observed in all the 3 Superclades. In particular (Figure 1), we notice that isolates from the USA are prevalently associated with Super-clade II, while isolates from Europe are prevalent in Super-clade III. Finally, Super-clade I incorporates the majority of viral isolates identified in Asia, and a more limited number of European isolates.

**Figure 1.**
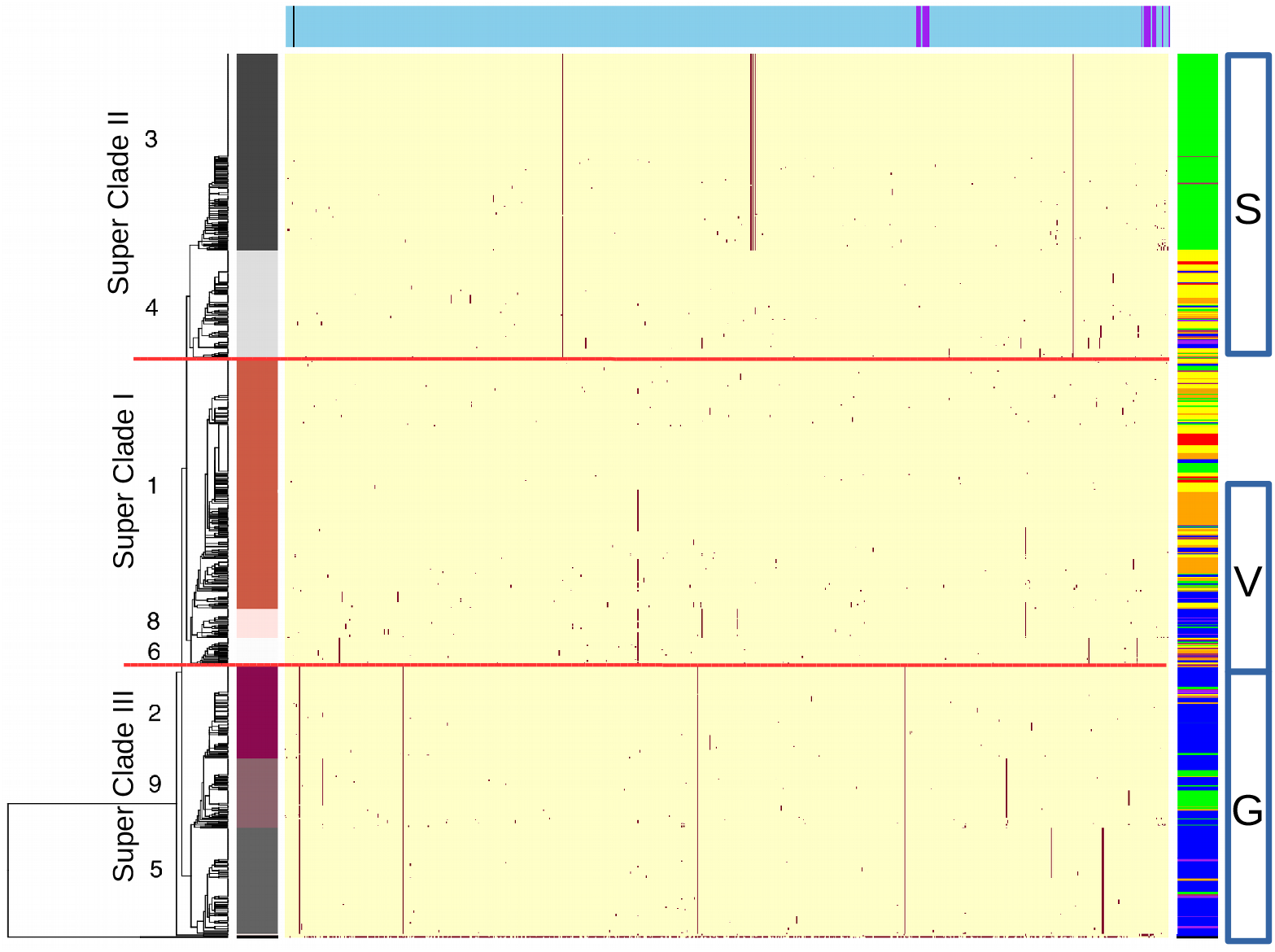
Heatmap of polymorphic sites in SARS-CoV-2 and SARS-CoV-2 like episomes. Color codes at the top of the heatmap indicate the type (light blue=substitution, black=deletion, purple=insertion) of the 844 genetic variants present in at least 2 distinct viral isolates. Genomic coordinates are represented on the X axis. The panels on the left indicate membership of one of the 9 clusters as defined in the text. Separation between the 3 Super-clades are indicated by red dotted lines. The panel on the right shows the geographic origin of the isolates (Green=USA, Yellow=China, not Wuhan, Red=China, Wuhan, Maroon=Australia, Blue=Europe, Orange=Asia, not China, Purple= South America, Black= non human host). The white boxes on the far right of the same panel represent the V, S and G clade as from the GISAID portal. Presence/absence of the polymorphic sites is indicated by a binary color code: Yellow=absent, that is the sequence is identical to the reference genome at that site, and Red=present.

Overall our analyses confirm the limited variability of the SARS-CoV-2 genome (Supplementary Figure S2), with an average number of 5.2 polymorphic sites observed between individual isolates and the reference genome. As reported in Supplementary Table S4, of the 983 sites that were polymorphic in at least 1 SARS-CoV-2 genome considered in the study, 310 (31.5%) were synonymous substitutions, 545 (55.4%) non-synonymous substitutions and 115 were associated with non-coding regions (5’ and 3’ UTR). Only a very limited number of indels were observed, with 3 single base insertions (2 in the 3’ UTR and 1 in the 5’ UTR) and 12 small insertions of which 10 were associated with the UTR regions of the genome.

The majority of polymorphic sites (703/983, 71.15%) are singletons associated with an individual viral isolate (Supplementary Figure S3). Interestingly, we notice that the average number of polymorphic sites in SARS-CoV-2 isolates are significantly (Wilcoxon p-value ≤ 1e-05) lower than the average variable sites between late-phase isolates of the SARS epidemic of 2003 (Supplementary Figure S2). Consistent with these reduced levels of variability, we observe that the 3 Super-clades identified in this study are defined by only a limited number of genetic polymorphic sites. For example, as evident from Figure 1, Super-clade II is characterized by the presence of only 2 polymorphic sites, which are common and specific to all the isolates of this group (8702 C->T and 28144 C->T). Super-clade III is defined by the presence of only 4 clade-specific variants (241 C->T, 3037 C->T, 14408 C->T and 23403 D->G). While no genetic variants are ubiquitous and exclusively associated with strains included in Super-clade I.

This notwithstanding we notice the presence of additional genomic variants, which could subtend the presence of smaller sub-clades.

Consistent with this observation, cluster stability metrics, based on the Dunn index, were strongly consistent with the presence of 9 clusters in the phenetic matrix of viral isolates. Clustering of viral isolates, based on the k-means algorithm with 9 clusters (Figure 1 and Supplementary Table S3) demonstrates a neat separation of viral episomes, with cluster number 7 corresponding to the two genome sequences of the SARS-CoV-2 like strains isolated from non human hosts. Cluster number 1 is formed by episomes that have limited variability with respect to the reference genome; cluster 2, 5 and 9 incorporate all the strains included in the European clade G; while clusters 3 and 4 correspond to the GISAID EpicoV clade S, with cluster 3 containing the majority of viral isolates prevalent in the U.S. Cluster 6 and cluster 8, which are composed of 35 and 40 viral strains, respectively, corresponds to a subset of the V clade as described in GISAID EpiCoV. While the majority of isolates included in cluster 8 have been isolated in Europe, cluster 6 does not seem to be associated with a specific geographic location. Notably, (Table 1 and Supplementary Table S4) we observe that the majority of polymorphic sites that discriminate between the 9 clusters identified by our analyses, are associated with nonsynonymous amino-acid substitutions, and several of these substitutions occur in viral genes that have been implicated in the modulation of the virulence of SARS and SARS-like associated coronavirus, including the spike protein and the RNA dependent RNA polymerase (nsp12) (Weiss and Navas-Martin, 2005). In order to test the possibility that these variations could reflect early hints of adaptive evolution of SARS-CoV-2 strains, evolutionary dynamics of viral super-clades were contrasted with patterns of evolution of the two closely related viral strains with different host specificity.

**Table 1.** List of polymorphic sites characteristic of each cluster of SARS-CoV-2 episomes. Position: indicates the genomic coordinate of the polymorphic site, Reference: the sequence of the reference genome and Alternative: the alternative allele, respectively. For variants associated with protein coding genes, the “AAresidue” and the “AAchange” columns are used to indicate the affected aminoacidic residue in protein coordinates and the predicted change in amino acid sequence, “-” indicates a silent substitution. The Gene column reports the corresponding geneor functionally annotated genomic element, while “Number of isolates” indicates the number of viral isolates in each cluster that have the polymorphism

| **position** | **reference** | **alternative** | **AAresidue** | **AA change** | **Gene** | **Number of isolates** |
| --- | --- | --- | --- | --- | --- | --- |
| *cluster1* |  |  |  |  |  |  |
| 11083 | G | T | 37 | L->F | nsp6 | 95 |
| *cluster2* |  |  |  |  |  |  |
| 241 | C | T | 81 | Na | 5'UTR | 125 |
| 3037 | C | T | 106 | - | nsp3 | 127 |
| 14408 | C | T | 323 | P->L | nsp12 | 125 |
| 23403 | A | G | 614 | D->G | spike | 127 |
| *cluster3* |  |  |  |  |  |  |
| 8782 | C | T | 76 | - | nsp4 | 246 |
| 17747 | C | T | 504 | P->L | nsp13 | 246 |
| 17858 | A | G | 541 | Y->C | nsp13 | 248 |
| 18060 | C | T | 7 | - | nsp14 | 247 |
| 28144 | T | C | 84 | L->S | orf8 | 248 |
| *cluster4* |  |  |  |  |  |  |
| 8782 | C | T | 76 | - | nsp4 | 137 |
| 28144 | T | C | 84 | L->S | orf8 | 138 |
| *cluster5* |  |  |  |  |  |  |
| 241 | C | T | 81 | Na | 5'UTR | 133 |
| 3037 | C | T | 106 | - | nsp3 | 133 |
| 14408 | C | T | 323 | P->L | nsp12 | 133 |
| 23403 | A | G | 614 | D->G | spike | 133 |
| 28881 | GGG | AAC | 203;204 | R->K,G->R | geneN | 133 |
| *cluster6* |  |  |  |  |  |  |
| 1397 | G | A | 198 | V->I | nsp2 | 35 |
| 11083 | G | T | 37 | L->F | nsp6 | 34 |
| 28688 | T | C | 139 | - | geneN | 35 |
| 29742 | G | T | 22 | Na | 3'UTR | 30 |
| *cluster8* |  |  |  |  |  |  |
| 11083 | G | T | 37 | L->F | nsp6 | 32 |
| 14805 | C | T | 455 | - | nsp12 | 40 |
| 17247 | T | C | 337 | - | nsp13 | 29 |
| 26144 | G | T | 251 | G->V | orf3A | 39 |
| *cluster9* |  |  |  |  |  |  |
| 241 | C | T | 81 | Na | 5'UTR | 77 |
| 1059 | C | T | 85 | T->I | nsp2 | 61 |
| 3037 | C | T | 106 | - | nsp3 | 78 |
| 14408 | C | T | 323 | P->L | nsp12 | 78 |
| 23403 | A | G | 614 | D->G | spike | 78 |
| 25563 | G | T | 57 | Q->H | orf3A | 78 |

As shown in Figure 2, comparison of intra-cluster variability, performed using only the 713 polymorphic sites that are associated with a single viral isolate, clearly demonstrates similar levels of variation between all the SARS-CoV-2 clusters identified in this study, with a slight (but not statistically significant, Wilcoxon test p-value 0.129) increase in variability for strains included in cluster 6.

**Figure 2.**
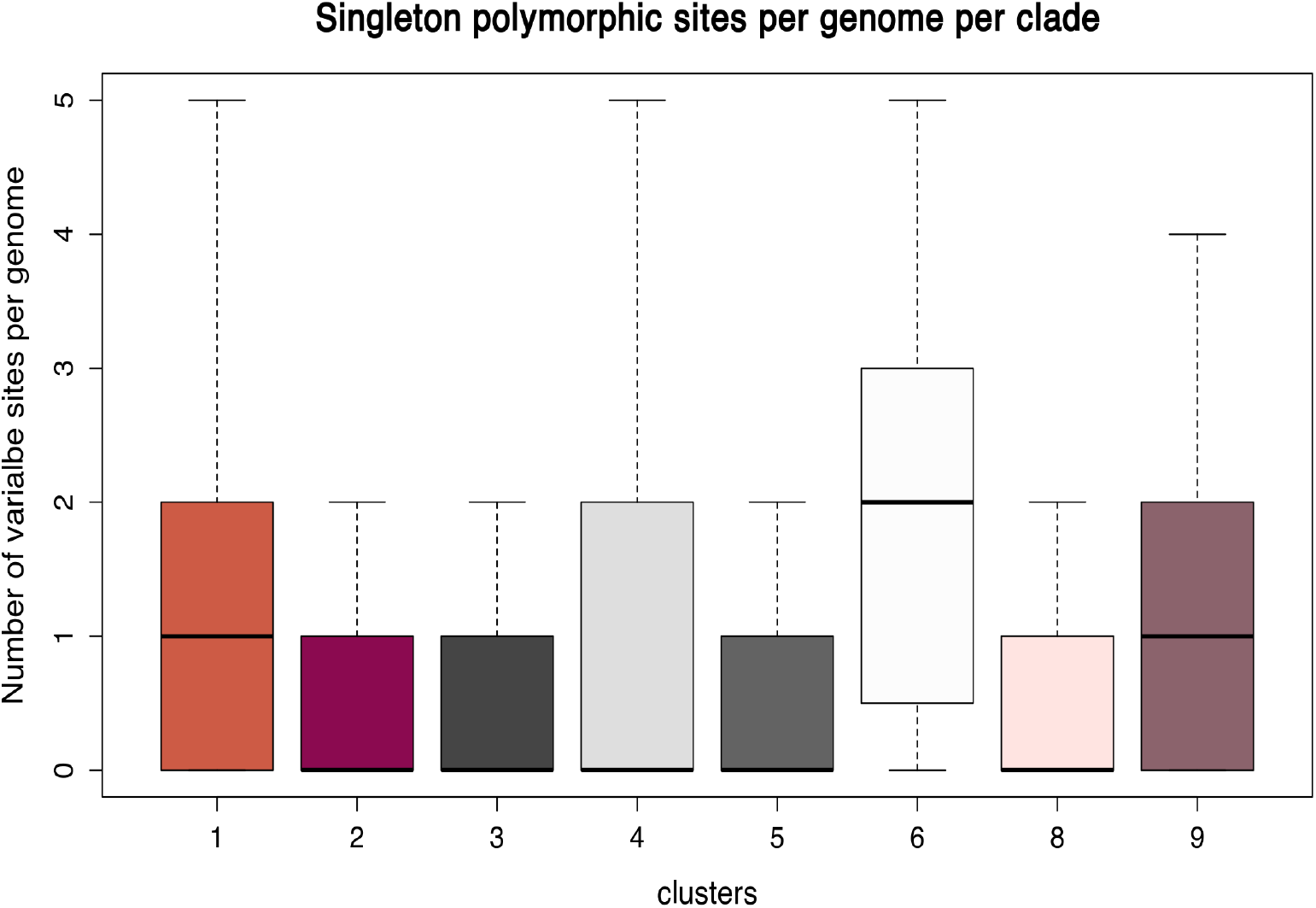
Boxplot of genomic variability for the 8 viral clusters of human-specific SARS-CoV-2 episomes, based on singleton polymorphic sites. Cluster color codes are as in Figure 1.

In order to identify regions of the genome which could be subject to distinct evolutionary pressures, plots of local genomic variability along the complete genomic sequence of SARS-CoV-2, were prepared by computing the proportion of polymorphic sites identified in each of the viral Super-clades on sliding genomic windows of 100 bp in size and overlapping by 50 bp. As shown in Figure 3 A-C, the observed patterns are remarkably similar between the 3 Super-clades, suggesting similar evolutionary dynamics. Moreover, we note that polymorphic sites are significantly enriched (Adjusted Fisher test p-value ≤1e-15 and ≤1e-12 respectively) in both the 5’ and 3’ UTR regions, while protein coding loci show considerably reduced variability.

**Figure 3.**
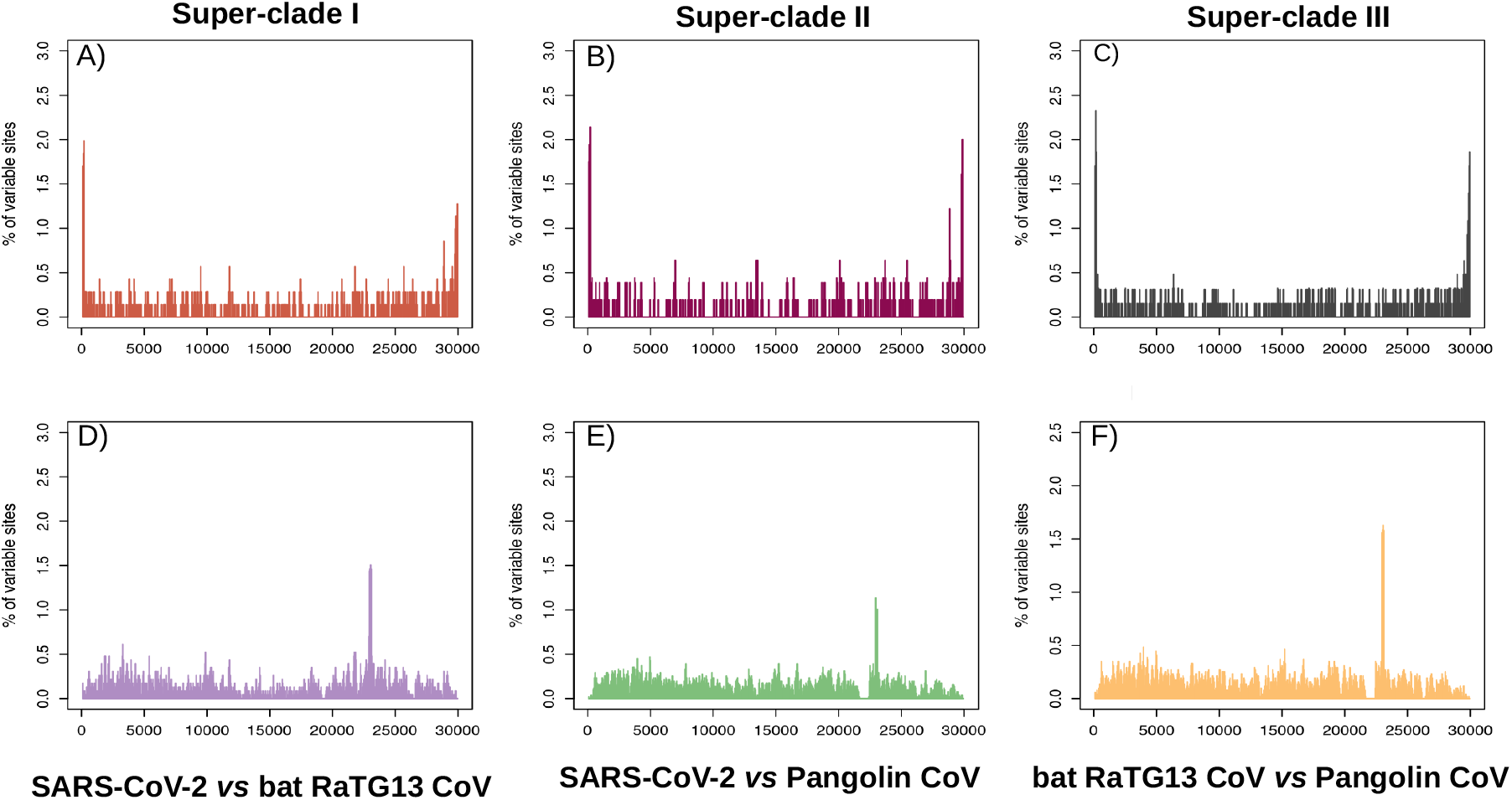
Plot of genomic variability calculated as the proportion of variable sites identified in overlapping genomic windows of 100 bp in:A) Super-clade I; B) Super-clade II; C) Super-clade III;D) the comparison of bat RaTG13 SARS-CoV-2 like against the reference SARS-CoV-2 genome; E) the comparison of Pangolin SARS-CoV-2 like against the reference SARS-CoV-2;F) the comparison of bat RaTG13 SARS-CoV-2 like against the pangolin SARS-CoV-2 like. Genomic coordinates are represented on the X axis, number of variable sites per window on the Y axis

Similar analyses performed by comparing the reference SARS-CoV-2 genome with the genomes of SARS-CoV-2-like coronavirus with a different host specificity, show the presence of a distinct pattern of variation in SARS-CoV-2-like genomes that is likely the result of adaptive evolutionary pressures (Figure 3 D-E). Indeed, hyper-variable genomic regions (Adjusted Fisher test p-value ≤1e-10 and ≤1e-5 respectively for comparisons of SARS-CoV-2 like CoVs isolated from bats and pangolins) between these genomes clearly correspond with protein coding loci, and in particular with the RDB protein domain of the S1 subunit of the spike protein - the domain which mediates the recognition of the host ACE2 receptor (see the peak at around position 23000 in Figure 3 D-E). A similar pattern is recovered also when genomes of bat and pangolin SARS-CoV-2 like CoVs are compared (Figure 3 F). Analyses of dN/dS patterns (Supplementary Table S5) indicate that, as expected, the RDB domain of the spike protein seems to be subject to strong diversifying selection between viral isolates with different host specificity (dN/dS > 1). On the contrary, in SARS-CoV-2 isolates we do not observe a dN/dS value > 1of the RDB domain, but we notice that, in all our comparisons, the gene coding for the spike protein shows the highest level of dN/dS among all protein coding genes, suggesting that - probably to facilitate adaptation to different hosts - this gene could be subject to relaxed selective pressure in coronaviruses.

Taken together, our analyses of variability profiles of SARS-CoV-2 and SARS-CoV-2-like coronaviruses are consistent with the notion that regions in the genome corresponding with increased variability, should be associated with relaxed purifying selection and/or increased diversifying selection. In this respect it is interesting to note that the 5’ and 3’ UTRs, which are the most variable regions of the genome between major SARS-CoV-2 strains, are among the least variable genomic regions, when SARS-CoV-2 and SARS-CoV-2 like strains with a different host specificity are compared.

## Discussion

Notwithstanding the limited variability of the genome, which prevents more detailed evolutionary analyses, our analyses provide no evidence for distinct evolutionary constraints acting on the different superclades and clusters of SARS-CoV-2 genomes. Although these results cannot be conclusive, the observation of similar dynamics of genomic variability, and that variable sites are prevalent at the 3’ and 5’ UTR genomic regions of SARS-CoV-2 indicate that it is unlikely that these differences should be the result of adaptive selection.

However, we notice that in SARS-CoV-2 the spike protein evolves under less constrained evolutionary dynamics compared to other genes. Indeed, in spite of the limited number of variable sites (135/3822) a dN/dS ratio of ~ 0.7 is observed for the spike gene, a value that is well above the value recovered for genes of similar size in the same genome and that would indicate that the spike gene is subject to weaker evolutionary constraints than the other protein coding genes. This possibly reflects a mechanism for the rapid adaptation to a more widespread range of hosts, as for example suggested by Menachery et al (Menachery et al, 2016).

Notwithstanding some limitations, our comparative analyses are consistent with the hypothesis that the biased geographic distribution, and the allelic differences observed between major viral SARS-CoV-2 clades are not the result of an adaptive evolutionary process, but are more consistent with founder effects on viral populations, coupled with the rapid spread of this novel virus.

Although our analyses do not suggest distinct evolutionary patterns, it remains unclear whether the genetic variants that discriminate between major viral clades could be related with differences in the virulence/pathogenicity of these clades. To address this issue it will be crucial to collect patient metadata, to sequence more genomes, and to enable the execution of retrospective statistical analyses.

## Materials and methods

The complete collection of high quality, complete SARS-CoV-2 genomes and associated metadata was accessed from the GISAID EpicoV (Shu and McCauley, 2017) platform on March 24th 2020. Genomes were aligned to the reference assembly of SARS-CoV-2 as available from Refseq (O’Leary et al 2016); Refseq accession NC_045512.2) by means of the nucmer (Marçais et al, 2018) program. Viral genomes of the SARS 2003 outbreak were retrieved from the NCBI virus database (Goodacre et al, 2018). Classification/association of strains to the 3 (early/middle/late) phases of the epidemic are according to Song et al 2005. Only isolates from the late phase of the epidemic were considered, based on considerations regarding the availability of a relatively high number of genomes (65) and the high level of similarity with the reference SARS-CoV genome.

Polymorphic sites were identified by using the show-snps utility of the nucmer package. Output files were processed by the means of a custom Perl script, and incorporated in a phenetic matrix, with variable positions on the rows and viral isolates in the columns. For all the isolates considered in the study, values of 1 were used to indicate presence of a variant, values of 0 its absence. dN/dS rates were computed on aligned CDS sequences using the Ka/Ks calculator tool (Zang et al, 2006) allowing for the selection of the most appropriate substitution model, based on the Akaike information criterion. The GY (Goldman and Yang,1994) model resulted to be the preferred model in all the settings herein tested. Only proteins longer than 100 amino acid residues and with more than 50 polymorphic sites in the CDS, were considered in this analysis. For SARS-CoV-2 this was limited to nsp12, nsp3, nsp4 and the spike protein.

Variability with respect to the reference NC_045512.2 SARS-CoV-2 assembly was computed on sliding windows of 100 bp, overlapped by 50 bp, by counting the proportion of variable genomic sites contained in each window (number of variable sites in the window, divided by the total number of variable sites), by using a custom Perl script. A Fisher-exact test, contrasting the local variability in a window with the average variability in the genome, was used to identify hypervariable regions. P-values were corrected using the Benjamini Hochberg procedure for the control of False Discovery Rate.

Functional effects of genetic variants as identified from genome alignments, were predicted by means of a custom Perl script, based on the annotation of the NC_045512.2 SARS-CoV-2 reference assembly.

Graphical representation of the data, basic statistical analyses and clustering of viral isolates were performed by means of the standard libraries of the R programming language. Similar to previous studies for the classification of SARS-CoV genomes (Chinese SARS Molecular Epidemiology Consortium, Song et al, 2005), clustering of viral genomes was performed by considering only sites that were polymorphic in at least 2 different genomes. Singleton sites, that is polymorphic sites observed only in one genome, were excluded from this analysis due to considerations regarding the limited information content, but also to the fact that, according to previous reports, these sites could be slightly enriched in sequencing errors (Song et al, 2005). Analogous considerations prompted us to execute the analysis of substitutions patterns both on the complete collection of polymorphic sites, and only on sites that were polymorphic in at least 2 genomes.

Determination of the optimal clustering solution was performed based on the Dunn Index metrics, as computed by the clValid R package (Brock et al, 2008).

## Acknowledgements

We thank ELIXIR Italy for providing the computing and bioinformatics facilities and Edward C. Holmes for his expert advice. We gratefully acknowledge the authors, originating and submitting laboratories of the sequences from GISAID’s EpiFlu™ Database on which this research is based. The list is detailed in Supplementary Table S1

## Supplementary Figures and Tables Legends

**Supplementary Table S1.** List of viral isolates included in the study. The table is in the same format as the submission acknowledgment table available from the GISAID EpiCoV website.

**Supplementary Table S2.** Rates of nucleotide substitution as identified from whole genome alignment. For every possible single nucleotide substitution (rows), the table reports the frequency of that substitution for: SARS-CoV-2_All vs SARS-CoV-2: alignments between SARS-CoV-2 genomes; SARS-Cov-2_2G vs SARS-CoV-2: as before, but computed only on polymorphic sites common to at least 2 genomes; SARS 2003L vs SARS 2003: alignment between all the SARS-CoV isolates from the late phase of the SARS 2003 epidemic (SARS 2003L) with the reference SARS-CoV genome; bat RaTG13 CoV vs SARS-CoV-2: alignment between the SARS-Cov-2 reference genome and the genome of a SARS-Cov-2 like of bat(RaTG13 genome); pangolin CoV vs SARS-CoV-2: alignment between the SARS-Cov-2 reference genome and the genome of a SARS-Cov-2 like pangolin

**Supplementary Table S3.** Cluster assignment of viral isolates. Accession number of the isolate in GISAID and the relative N° of polymorphic sites with respect to the reference genome, are also reported

**Supplementary Table S4.** Functional annotation of polymorphic sites. The table lists all the 983 polymorphic sites identified from the comparison of 1113 SARS-Cov-2 genomes with the Refseq assembly. “Pos” reports the genomic coordinates of the polymorphic site, followed by the sequence on the reference assembly (Ref) and the alternative sequence (Alt), with “.” indicating insertion/deletions. For variants associated with protein coding genes, “AA pos” and “AA change” are used to indicate the affected amino-acidic residue in protein coordinates, and the predicted change in amino acid sequence, respectively. Type indicates the predicted functional effect: “S” indicates a silent substitution, “NS” a nonsynonymous substitution, “FS” a frameshift, “Non-coding”, that the variant is associated with a non protein coding region of the genome, “Na”, stands for not applicable. “Functional element” reports the corresponding gene or functionally annotated genomic element, and “N° of genomes” indicates the number of viral isolates in each cluster that have the polymorphic site. The equivalent information for every cluster (number of isolates that have the variant) is reported in the columns “Cluster 1” to “Cluster 9”, for clusters from 1 to 9 respectively.

**Supplementary Table S5.** dN/dS ratio. The table reports the dN/dS ratio, computed by the means of the KaKs_Calculator for all the protein coding genes longer than 100 aa residues and with more than 50 variable sites associated with their CDS. Proteins are indicated by their gene symbol on the rows. “spike_rbd” indicates the spike recognition binding domain, “concat_no_spike”, indicates a supergene formed by all the protein coding genes in the genome with the exclusion of the spike protein. “Na” stands for not applicable. bat RaTG13 CoV vs SARS-CoV-2: dN/dS between SARS-CoV-2 and the bat RaTG13 CoV; pangolin CoV vs SARS-CoV-2: dN/dS between SARS-CoV-2 and the SARS-CoV-2 like CoV isolated from pangolin specimens; SARS-CoV-2: dN/dS for SARS-CoV-2.

**Supplementary Figure S1.** Heatmap of nucleotide substitution frequencies. The heatmap displays nucleotide substitution frequencies, as derived from whole genome alignment as reported in Table S1. Frequencies are reported in each cell of the heatmap. A gray (low) to blue (high) gradient of color is used. The suffix “_AllG” and “_2G” are used to indicate substitution profiles derived from the analysis of all the variable sites (“_AllG”) or of only sites variable in at least 2 genomes (“_2G”).

**Supplementary Figure S2.** Histogram of the number of variable sites identified in: A) any SARS-CoV-2 genomes included in this study, with respect to the reference SARS-CoV-2 genome; B) SARS-CoV genomes from the late phase of the 2003 epidemic with respect to the SARS-CoV reference genome.

**Supplementary Figure S3.** Prevalence of polymorphic sites in viral isolates. Number of viral genomes is reported on the X axis. Log scaled counts of the number of polymorphic sites supported by that number of genomic sequences is indicated on the Y axis

