## Supplementary figures and images for "Comparative genomics suggests limited variability and similar evolutionary patterns between major clades of SARS-CoV-2"

### Supplementary Figures

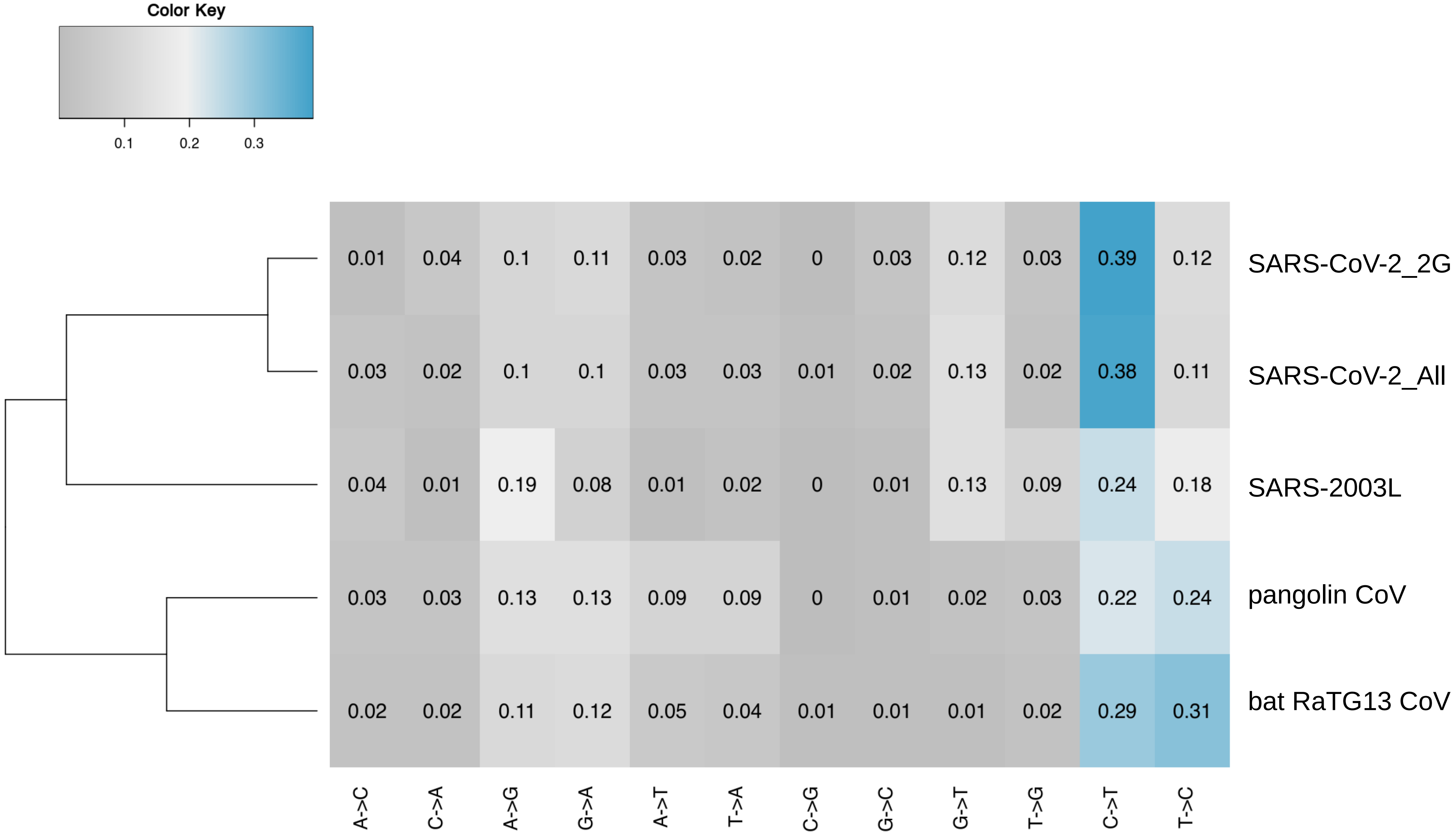

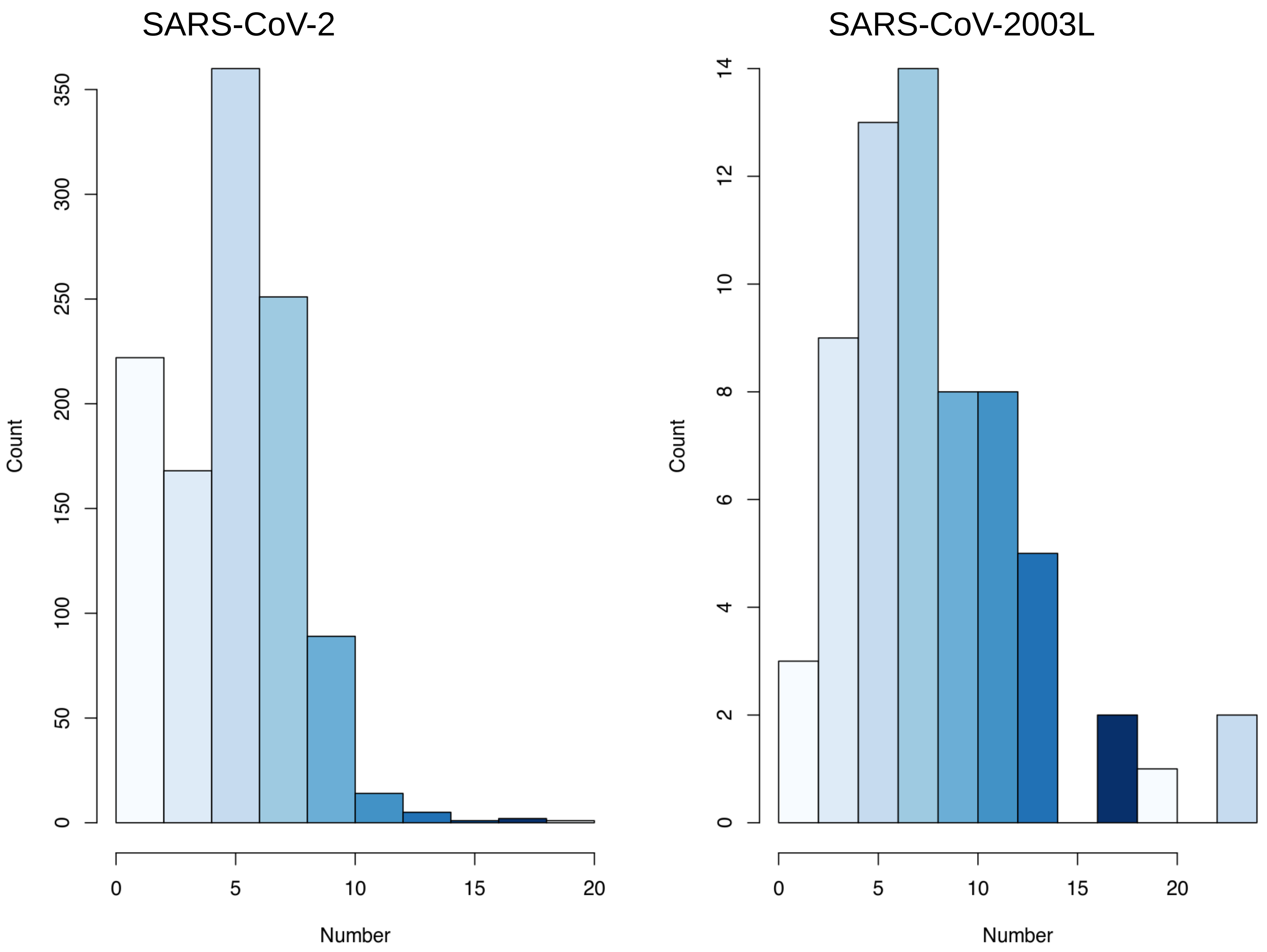

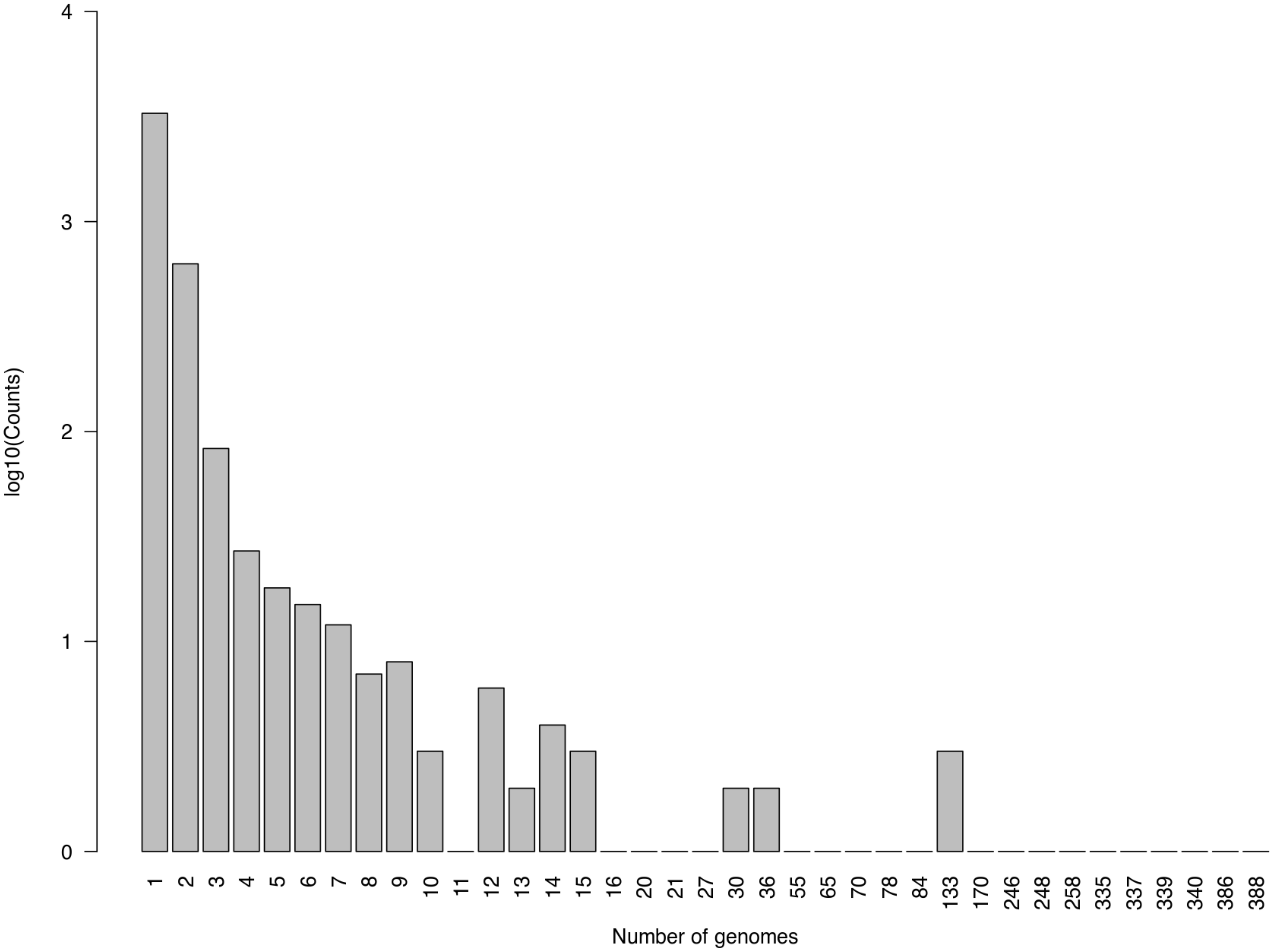
